# A manual multiplex immunofluorescence method for investigating neurodegenerative diseases

**DOI:** 10.1101/533547

**Authors:** Alexander J. Ehrenberg, Dulce Ovando Morales, Jorge Santos Tejedor, Mihovil Mladinov, Antonia Piergies, Jan Mulder, Lea T. Grinberg

## Abstract

**BACKGROUND:** Neurodegenerative diseases feature stereotypical deposits of protein aggregates that selectively accumulate in vulnerable cells. The ability to simultaneously localize multiple targets in situ would facilitate discovery and validation of molecular pathways underlying neurodegenerative diseases. Immunostaining methods offer in situ detection of targets. Most proposed protocols for multiplexing immunostaining require sophisticated equipment not available for standard labs. Effective stripping of antibodies, allowing successive rounds of staining while maintaining tissue and antigen integrity, is the main roadblock for enabling multiplex immunostaining in standard labs. Antibody stripping techniques have not been fully validated for neurodegenerative disease research. Moreover, despite the increased complexity of multiplex immunostaining protocols, quality control steps have not been regularly implemented.

**NEW METHOD:** Aiming to create protocols for multiplex localization of neurodegenerative-related processes, without the need for specialized equipment, we evaluated several antibody stripping techniques. We also recommend quality control steps to validate stripping efficacy and ameliorate concerns of cross-reactivity and false positives.

**RESULTS:** The proposed protocol using β-mercaptoethanol enables reliable antibody stripping across multiple rounds of staining and minimizes the odds of cross-reactivity while preserving tissue and antigen integrity.

**COMPARISON WITH EXISTING METHODS:** Our proposed method accounts for the intricacies of suboptimal human post-mortem tissue and the need to strip markers bound to highly aggregated proteins. Additionally, it incorporates quality control steps to validate antibody stripping.

**CONCLUSIONS:** Multiplex immunofluorescence methods for studying neurodegenerative diseases in human postmortem tissue are feasible even in standard laboratories. Nevertheless, evaluation of stripping parameters in optimization and validation phases of experiments is prudent.

## 1. Introduction

High-throughput genomic and proteomic methods have enabled probing of the milieu of molecular biology underlying complex diseases. However, these methods destroy tissue integrity and thus, results require validation in diseases such as neurodegenerative diseases in which different cell populations show distinct levels of vulnerability and changes (Fu et al., 2018; Seeley, 2008; Seeley et al., 2009). Recent developments in single-cell -omics methods allow more granular probing of the selective changes in individual cells. Unfortunately, because single-cell methods inevitably require a tissue dissociation step that compromises cellular membranes, most of these methods rely on cell nuclear rather than cytoplasmic composition, whereas most protein inclusions associated with neurodegenerative disease localize to the cytoplasm or in the extracellular space, thus it makes it challenging to distinguish affected versus unaffected cells.

Immunostaining remains an attractive validating method to investigate the relevance of results disclosed by these high-throughput techniques because it can unlock insights into molecular cascades while preserving tissue integrity providing a spatial overview with subcellular resolution. However, potentially valuable information often gets locked up in a tissue section due to the inherent low-throughput nature of these methods. Methodological constraints on classical immunostaining methods restrict probing to a few markers in each tissue section. Traditional immunohistochemistry and immunofluorescent methods rely on visualizing primary antibody binding sites using a detection system containing a secondary antibody against the species in which the primary antibody was raised. To avoid cross-reaction between conspecific antibodies, some protocols suggest direct fluorescent conjugation of primary antibodies. The advent of tyramide signal amplification (TSA) improved the options for combining conspecific antibodies in the same sample while providing signal amplification if combined with an antibody stripping technique. (Buchwalow et al., 2018; Chao et al., 1996; Dixon et al., 2015; Lim et al., 2018; Mansfield, 2017; Pirici et al., 2009; Roy et al., 2019; Sorrelle et al., 2019; Stack et al., 2014; Wang et al., 1999; Zhang et al., 2017). In short, TSA uses a peroxidase-mediated reaction to covalently deposit fluorophores or chromogens to tyrosine side chains proximal to the target epitope (Lim et al., 2018; Wang et al., 1999). Because of the covalent bond, stripping primary and secondary antibodies based on protein denaturing will not affect tissue bound tyramide-fluorophores.

Nevertheless, even if issues surrounding the simultaneous use of conspecific antibodies are ameliorated, multiplex immunostaining protocols are still limited by the number of spectrally separated detection channels. Signal of chromogenic reporters such as 3,3’-Diaminobenzidine detected by brightfield microscopy overlap and limits detection to two or, rarely, three probes simultaneously especially when targets are co-existing in the same cell types (Dixon et al., 2015; Gown et al., 1986; Ilie et al., 2018; Lan et al., 1995; Nakane, 1968; Tramu et al., 1978). Fluorescence microscopy offers the possibility to detect more probes simultaneously by utilizing molecules with specific excitation and emission spectra as reporters, allowing selective visualization of different probes in separate channels which are subsequently overlapped digitally (Coons, 1961; Coons et al., 1941; Coons et al., 1942; Coons and Kaplan, 1950). Still, limitations in a camera’s detectable wavelengths and spectral overlap between fluorophores, most fluorescent microscopes are still limited to detecting up to five markers. More recently, alternative methods for in situ detection were proposed, such as those replacing fluorescent-based reporters for quantum dots that are directly conjugated in primary antibodies. Quantum dots allow for a higher number of probes to be detected at the same tissue because of reduced spectral overlap (Byers and Hitchman, 2011; Chan et al., 2005; Chen et al., 2013; Dixon et al., 2015; Krenacs et al., 2010; Mansfield, 2017; Prost et al., 2016; Sweeney et al., 2008; Tholouli et al., 2008; Wu et al., 2003; Xu et al., 2013), however, quantum dots and other similar techniques require expensive, specialized equipment.

Thus, a fundamental roadblock precluding multiplex immunostaining in standard laboratory settings is the effective stripping of antibodies, especially since many antibodies use for clinical diagnostics are mouse monoclonal antibodies. Antibody stripping enables the use of conspecific antibodies and multiple rounds of staining, imaging, and stripping, thus increasing the number of markers probed in the same histological section. Several methods for stripping antibodies have been proposed, but almost none have been optimized to strip antibodies bond to highly aggregated proteins, as it is the case for those employed in neurodegenerative disease research. Further, most methods tend to affect tissue integrity, precluding additional rounds of staining. Here, we evaluate different multiplex histology protocols to identify a reliable, low-cost multiplex immunostaining method for routine use in the setting of a standard neuropathology laboratory. Conceptually, our technique of choice would result in a complete, efficient, and reliable stripping of a broad array of antibodies while preserving tissue integrity and antigenicity in FFPE samples, across multiple rounds of stripping and staining. We chose to use fluorescence detection because immunofluorescence microscopes are widely available and accommodate more detection channels than brightfield microscopy. Next, we make recommendations on limitations, appropriate use, and necessary steps to incorporate techniques into experimental setups.

## 2. Material and Methods

Towards establishing our protocol, we first created a list of principles and established approaches to elute antibodies based on a literature survey. Next, we implemented these in our standard immunofluorescence protocols in two experimental stages. In the first stage, we tested the most promising techniques to strip two antibodies used to label neuronal cell types, and most importantly, one antibody used to label phospho-Ser202-tau (phospho-tau), a common marker in Alzheimer’s disease research. Phospho-tau positive intracellular deposits include highly aggregated inclusions and are commonly found in neurodegenerative research. In the second stage, we tested the best stripping method, according to the results of stage 1, across five rounds of staining and stripping using an expanded panel of six antibodies (Figure 1). Here, we evaluated whether the stripping method could be applied to a larger number of antibodies while preserving tissue and antigen integrity. Multiple rounds of staining are crucial in multiplex protocols relying on immunofluorescence because every single round is limited to three to five channels. In the same line, the signal of at least one marker has to remain stable throughout the multiple rounds to allow digital co-registration of images from different rounds.

**Figure 1:**
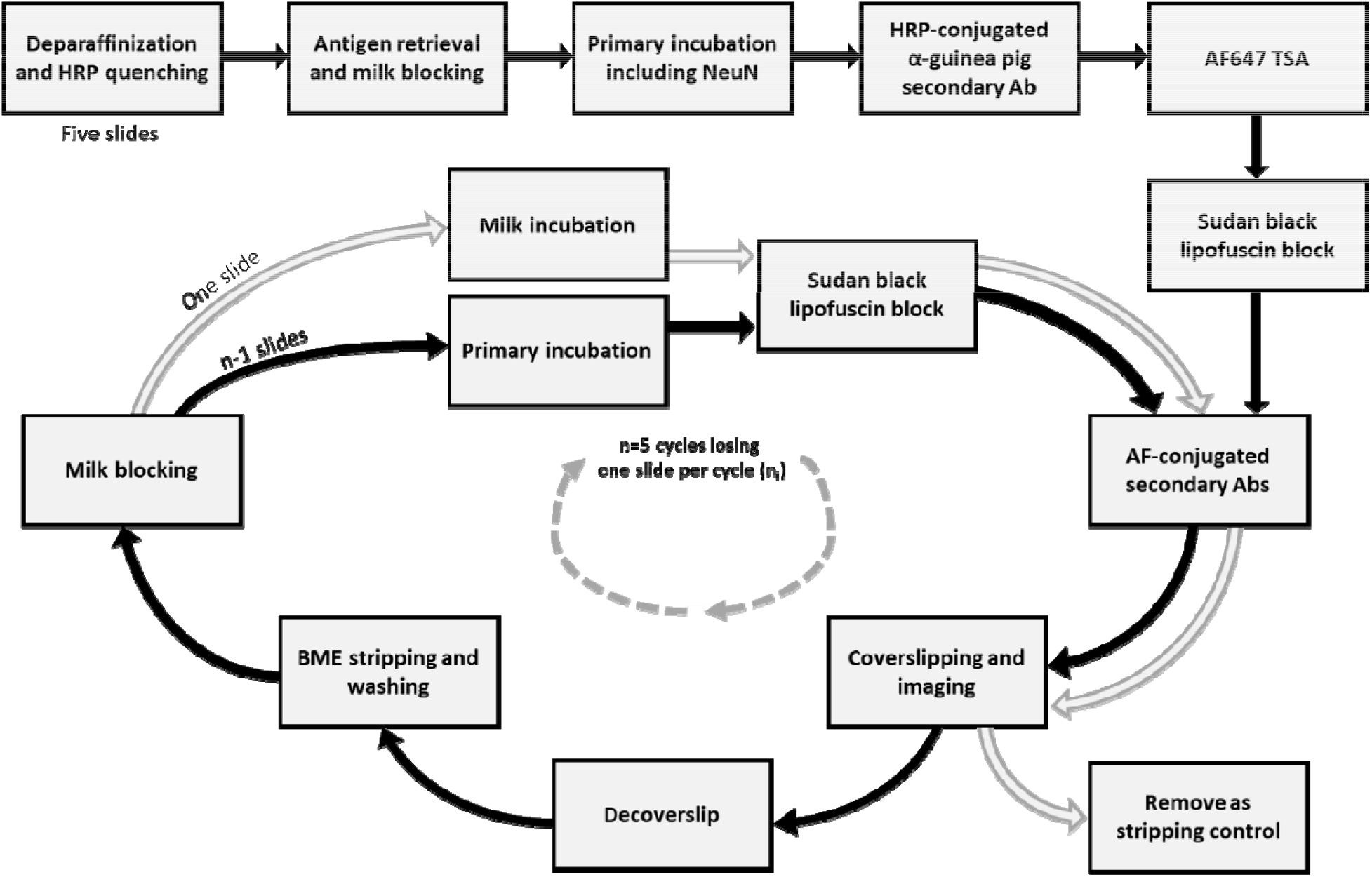
Flowchart depiction of tissue and antigen preservation experiment with each antibody set starting with five slides. One slide is removed each round for stripping checks and then kept in storage.

### 2.1 Literature review and selection of candidate stripping techniques

We searched PubMed for the terms “antibody elution histology” and “multiplex immunohistochemistry.” “Antibody elution histology” yielded 462 results and “multiplex immunohistochemistry” yielded 2021 results. In total, these combined searches yielded 2480 unique results ranging from November 1964 to May 2019. These results were manually filtered to exclude articles unrelated to histology. From the remaining, we identified seven main stripping strategies previously used in multiplex immunohistochemistry protocols. Although stripping techniques may either elute or denaturate antibodies, both terms are used interchangeably in the literature. These seven strategies are: (1) heat-based, particularly common in multiplex protocols using TSA (Ilie et al., 2018; Jufas et al., 2008; Lan et al., 1995; Lim et al., 2018; Mansfield, 2017; Parra et al., 2017; Roy et al., 2019; Saylor et al., 2018; Sorrelle et al., 2019; Stack et al., 2014; Toth and Mezey, 2007; Wegner et al., 2017; Zhang et al., 2017); (2) a glycine/SDS solution at a low pH (Bolognesi et al., 2017; Gut et al., 2018; Nakane, 1968; Narhi et al., 1997a; Pirici et al., 2009; Sorrelle et al., 2019); (3) commercially available denaturing solutions for IHC with proprietary formulations (Buchwalow et al., 2018); (4) β-mercaptoethanol (BME) denaturation (Bolognesi et al., 2017; Gendusa et al., 2014; Kim et al., 2012; Mansfield, 2017; van den Brand et al., 2014) which reduces the disulfide bonds present in antibodies, thus breaking down their tertiary structure (Capel et al., 1980; Crivianu-Gaita et al., 2015); (5) a low-pH oxidizing solution of KMnO_4_ and H_2_SO_4_ (Glass et al., 2009; Tramu et al., 1978); (6) chaotropic salts (Bolognesi et al., 2017; Gut et al., 2018; Narhi et al., 1997a; Narhi et al., 1997b); (7) combining antibodies with drastically different abundances (Wang et al., 1999).

From these seven candidate principles, we initially excluded three. Techniques based on a low-pH oxidizing solution (Glass et al., 2009; Tramu et al., 1978) tend to inactivate fluorophores and damage the tissue. Chaotropic salts, such as guanidinium, are not effective at stripping antibodies bound to highly aggregated, spatially organized targets and requires steps to recover antigen conformations following the denaturation step (Bolognesi et al., 2017). Finally, the technique that relies on making the *a priori* assumption that some targets are present in less abundance than others (Wang et al., 1999) is extremely limited in nature and based on implicit assumptions regarding the sensitivity of microscope cameras and variation in biological samples. Table 1 depicts the four stripping techniques tested in Stage 1.

**Table 1:** Candidate stripping techniques tested for Stage 1.

| Strategy | Description | Reference for protocol |
| --- | --- | --- |
| Heat | Microwave in citrate buffer with 0.05% tween. High setting until boiling then 15 minutes at 20% (Note: referenced paper suggests 5 minutes at 50%). | Toth and Mezey (2007) |
| Glycine/SDS at low pH | Glycine mixed with SDS, at pH 2.0 and heated to 50°C for 30 minutes. | Pirici et al. (2009) |
| Denaturing solution | One part of solution A to two parts of solution B from the Denaturing Solution Kit from Biocare Medical | Manufacturer instructions (part DNS001L) |
| BME | Tris-HCl mixed with SDS and BME, heated to 56C and incubated for 30 minutes. Washed in TBS. | Gendusa et al. (2014) |

### 2.2 Experimental Stage 1: choosing the best stripping technique

#### 2.2.1 Rationale

In this stage, we tested stripping efficiency and effects on immunogenicity for each of the four candidate stripping techniques using three different primary antibodies representing a protein aggregate in Alzheimer’s disease and two cell-type markers, all three from different species. Each primary antibody was tested in four serial histological slides, one for each stripping technique. We incubated the slides with one of the three primary antibodies, followed by incubation with a host species-specific HRP-conjugated secondary antibody developed with TSA using a tyramide-conjugated to Alexa Fluor 647 (AF647). This step confirms proper antigen retrieval, and the presence and distribution of the target. Because TSA covalently deposits fluorophores to the tissue, the fluorescent signal should remain even after the primary and secondary antibodies are stripped (Lim et al., 2018; Wang et al., 1999). For this reason, AF647 signal represents the distribution of antibody reactivity independent of the stripping here. Thus, the signal in this channel (Cy5) is set as the standard. Next, we applied each stripping technique. To test whether the primary antibodies were successfully stripped, we incubated the same slides again with same species-specific HRP-conjugated secondary used before, but this time, developed the reaction with tyramide-conjugated with Alexa Fluor 546 (AF546). If the stripping was successful, the reaction would yield no signal in the DsRed channel. Presence of any signal in the DsRed channel would indicate a failure in the stripping. Finally, to test whether the target antigens remained preserved after the stripping treatment, we re-incubated the same slides with the same primary antibodies followed by HRP-conjugated secondary antibodies and developed the reaction with tyramide conjugated to Alexa Fluor 488 (AF488). Preservation of the antigen of interest should yield nearly complete signal overlap between AF488 (visualized in the GFP channel) and AF647 (visualized in the Cy5 channel).

#### 2.2.2 Tissue source

De-identified human brain tissue was supplied by the Neurodegenerative Disease Brain Bank from the University of California, San Francisco’s Memory and Aging Center. Post-mortem interval of the cases varied from 7 to 30 hours. Brain samples were fixed in 10% neutral buffered formalin for 72 hours and transferred to PBS-azide at 4°C for long-term storage. 8μm thick FFPE sections were mounted on Ultra Bond adhesive slides (SL6023-1, Avantik BioGroup) and incubated in an oven at 65°C for at least 18 hours before further processing. The collection of these tissues and use in the current study was approved by the institutional review boards from the University of California, San Francisco and this experiment was considered to belong to the non-human category (postmortem and de-identified) by the same IRB.

#### 2.2.3 Primary antibodies

We chose a commonly used monoclonal antibody that targets pathologic aggregates of hyperphosphorylated tau protein (CP13, phospho-Ser202 tau, 1:800; gift of Peter Davies, New York). We also used antibodies against Fox-3 (RBFOX3 NeuN, 1:600; #266004, Synaptic Systems, Goettingen, Germany) and glial fibrillary acidic protein (GFAP, 1:1800; #ab68428, Abcam) that label neurons and astroglia, respectively. Table 2 depicts the host species, vendor information, and dilution used for all primary antibodies used in this study, including those used in Experimental Stage 2.

**Table 2:** Primary antibodies used in this study

| Antibody | Host | Label | Source | Working dilution |
| --- | --- | --- | --- | --- |
| CP13 | Ms, monoclonal IgG<sub>2b</sub> | Phospho-tau (Ser 202) | Gift from Peter Davies, New York, NY | 1:800 |
| TMEM119 | Rb, polyclonal | Yolk-sac derived microglia | Sigma Aldrich (HPA051870) | 1:500 |
| PHF-1 | Ms, monoclonal IgG<sub>1</sub> | Phospho-tau (Ser 396/404) | Gift from Peter Davies, New York, NY | 1:9000 |
| GFAP | Rb, monoclonal EPR1034Y | astroglia | Abcam (ab68428) | 1:1800 |
| MC1 | Ms, monoclonal IgG<sub>1</sub> | Conformational tau | Gift from Peter Davies, New York, NY | 1:1000 |
| T18 | Rb, polyclonal | Oligomeric tau | Rakez Kayed, Galveston, TX | 1:1500 |
| NeuN | GP, polyclonal | Neurons | Synaptic Systems (266 004) | 1:600 |

#### 2.2.4 Immunostaining protocol

Except for the stripping step, all steps of the immunostaining protocol are routine in histology labs. Slides underwent serial deparaffinization steps as follows: three ten-minute immersions in xylene, two two-minute immersions in 100% ethanol, two two-minute immersions in 96% ethanol, and one two-minute immersion in 80% ethanol. Slides were then immersed in a solution of 3% hydrogen peroxide (H_2_O_2_) and 80% methanol for 30 minutes to quench any endogenous peroxidase. Sections underwent three two-minute washes in distilled water (dH_2_O) before being transferred into a 10% solution of 0.1M citrate buffer pH 6.0 with 0.05% tween-20 for antigen retrieval. In the antigen retrieval solution, sections were cycled through an autoclave set at 121°C for five-minutes. Following antigen retrieval, sections were left to cool to room temperature (RT) for approximately 60 minutes. After thorough washing in a solution of 1x PBS and 0.05% tween (PBST), sections were immersed in a solution of 5% milk with 0.05% tween (herein referred to as milk) for 30 minutes to block unspecific binding. Sections were then incubated in CP13 (1:800, gift of Peter Davies), NeuN (1:600; #266004, Synaptic Systems), or GFAP (1:1800; #ab68428, Abcam) diluted in 5% milk PBS solution for 16 hours overnight at RT.

After washes in PBST, sections were incubated in a 1:400 concentration of species-specific HRP-conjugated secondary antibody (goat-α-mouse IgG(H+L)-R-05071, Advansta; goat-anti-guinea pig IgG (H+L) – (R-05076, Advansta, or goat-anti-Rabbit IgG (H+L)-R-05072, Advansta) diluted in PBST for 60 minutes. Following additional PBST washes, the antibodies were developed with TSA following the manufacturer’s instructions with a solution of 1:100 Alexa Fluor 647 (AF647) Tyramide (B40958, Thermo Fisher) and 1:100 of 100x H_2_O_2_ in 1x tris-buffered saline for 15 minutes and was followed by PBST washes. Following TSA, one of several possible stripping techniques was employed according to the protocols given by the references noted in Table 1.

Next, sections were re-incubated in a 1:400 species-concentration of specific HRP-conjugated secondary antibody for 60 minutes, followed by PBST washes, and an additional TSA step with Alexa Fluor 546 (AF546) Tyramide (B40954, Thermo Fisher) for 15 minutes. The TSA step was ended with a PBST wash. Finally, the sections were re-incubated with the original primary and secondary antibodies as previously described, followed by an additional TSA step according to the manufacturer’s instructions with a solution of 1:100 Alexa Fluor 488 (AF488) Tyramide (B40953, Thermo Fisher) and 1:100 of 100x H_2_O_2_ in 1x tris-buffered saline. This amplification occurred for 15 minutes and was followed by PBST washes.

After a transfer through 70% ethanol, the sections were treated with a solution of 0.8% Sudan Black-B in 70% ethanol for 35 minutes for blocking lipofuscin autofluorescence, followed by two ten-second washes in 70% ethanol to remove excess Sudan black-B. Sections were re-hydrated in PBS then coverslipped with Prolong Glass Antifade Mountant with NucBlue (P36981, Thermo Fisher).

#### 2.2.5 Image acquisition

Slides were imaged on a Zeiss AxioImager.A2 microscope equipped with a Zeiss Colibri 7:Type FR-R[G/Y]CBV-UC 7-channel fluorescence light source. NucBlue (Hoechst 33342) was visualized with a DAPI filter set, AF488 visualized with a GFP filter set, AF546 with a DsRed filter set, and AF647 with a Cy5 filter set.

### 2.3 Stage 2: Evaluation of tissue and antigen preservation through multiple rounds

#### 2.3.1 Rationale

β-mercaptoethanol-based (BME) stripping was the only technique that successfully stripped all antibodies and did not cause dissociation of the tissue from the slide (Figs. 2–4). In this Stage, we interrogated whether BME-based stripping could accommodate up to five rounds of staining, coverslipping, imaging, decoverslipping, stripping, and restaining in six antibodies. We combined the primary antibodies in groups of three and stained five slides per group. Importantly, all antibody combinations included NeuN, that was selected as an optimal marker to allow co-registration of images obtained across different rounds of staining. Thus, NeuN was developed with TSA on round one only, as the corresponding fluorescent signal should remain throughout the multiple rounds despite stripping of antibodies. The other two antibodies in each group were always developed with species-specific secondary antibodies conjugated with AF488 or AF546 to allow removing fluorescent signal together with antibody stripping.

**Figure 2:**
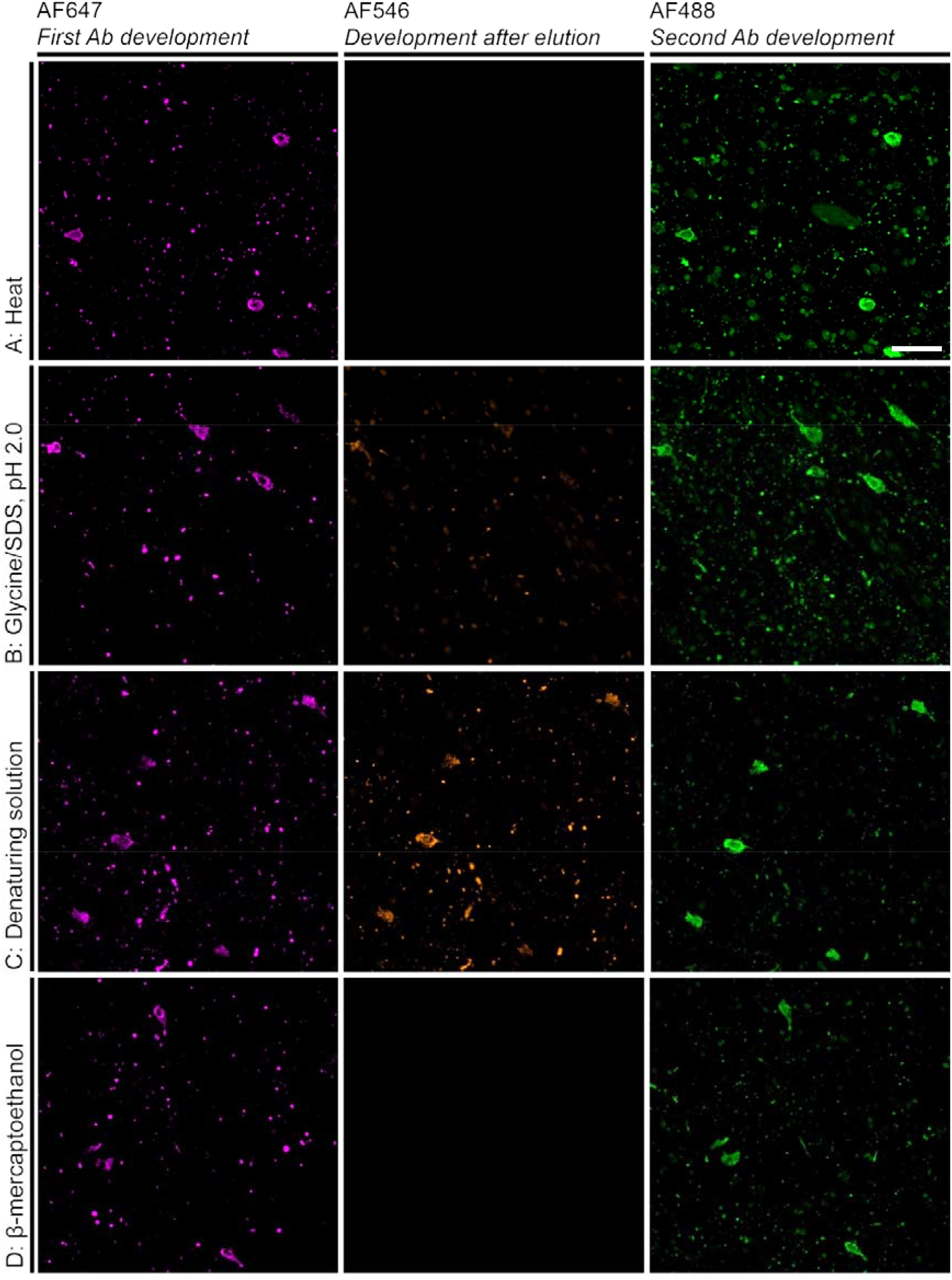
Photomicrographs of the validation experiments for CP13. Successful stripping and antigen preservation are represented by overlapping signal from AF647 and AF488 with zero signal from AF546 (Heat and β-mercaptoethanol, A and D). The Glycine/SDS, pH 2.0 (B) resulted in reduced, but noticeable signal from AF546, and denaturing solution (C) resulted in significant signal from AF546. Scale bar: 10 μm

**Figure 3:**
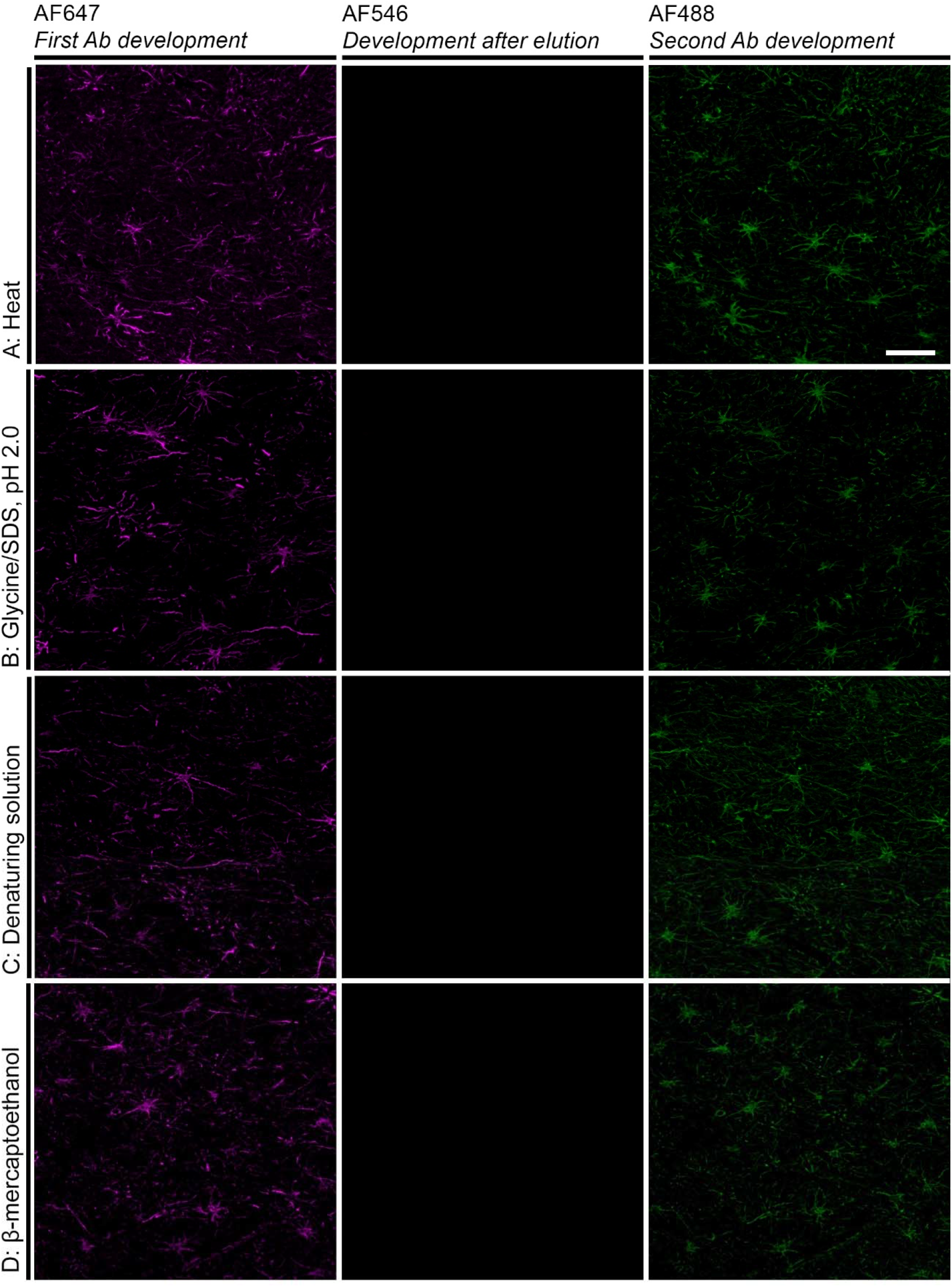
Photomicrographs of the validation experiments for GFAP. Successful stripping and antigen preservation is represented by overlapping signal from AF647 and AF488 with zero signal from AF546 for all tested stripping strategies.. Scale bar: 10 μm

**Figure 4:**
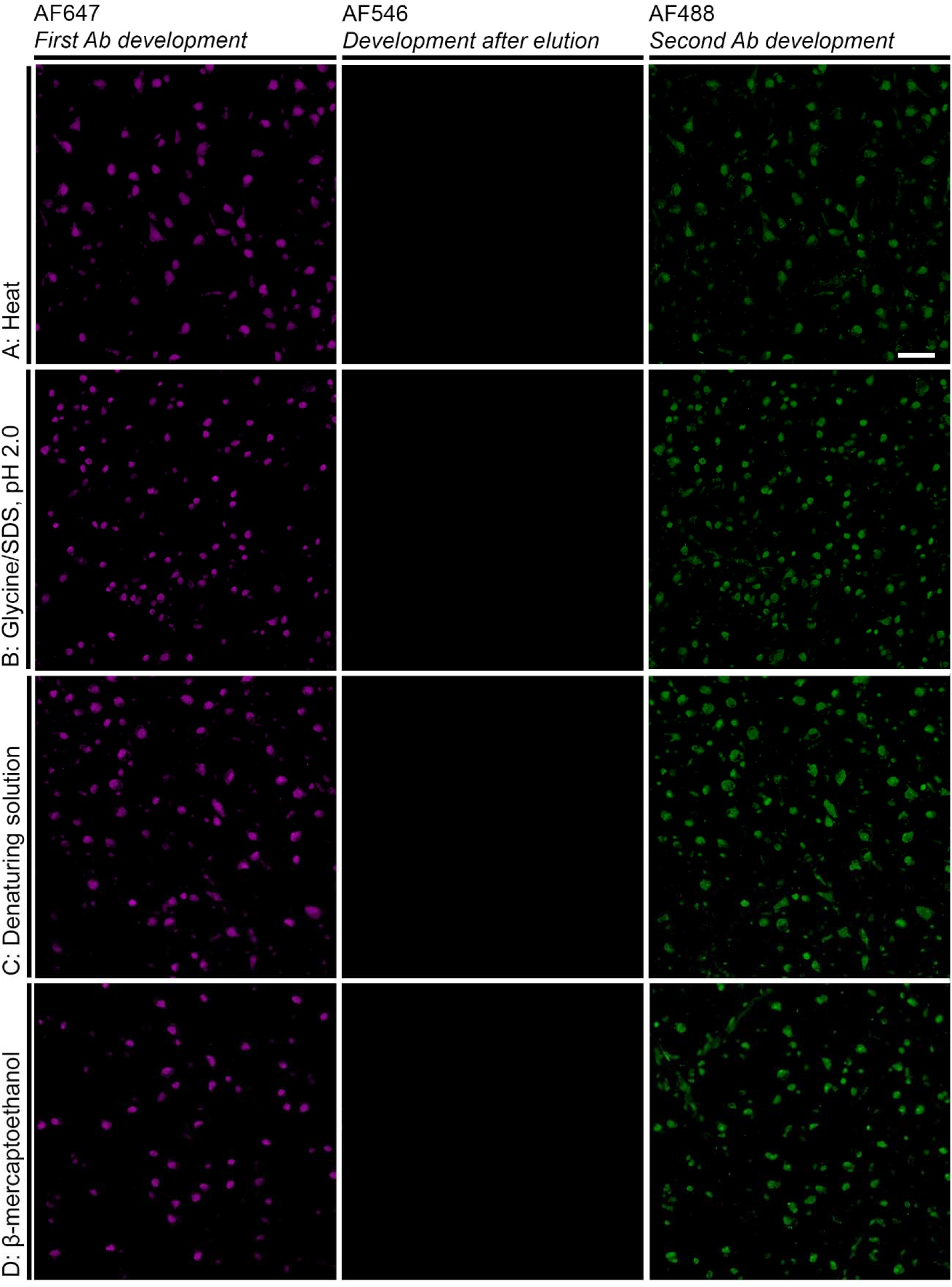
Photomicrographs of the validation experiments for NeuN. Successful stripping and antigen preservation is represented by overlapping signal from AF647 and AF488 with zero signal from AF546 for all tested stripping strategies. Scale bar: 10 μm

In the first round of staining, all five slides of each set underwent the same procedure. In the subsequent rounds, one of the slides of each set was set aside and incubated with 5% milk PBS instead of the primary antibodies. Thus, on rounds 2, 3, 4 and 5 only 4, 3, 2 and 1 slides of the set underwent primary antibodies incubation (Fig. 1). Thus we could test the stripping efficiency after each round. For each set, we tested the maximum number of rounds accommodated by the protocol (up to 5), determining at which round, if any, the tissue started to dissociate from the slide and/or the signal for a given antibody was markedly decreased compared to round 1.

#### 2.3.1 Tissue selection and primary antibodies

We tested this protocol successfully in different cases. For this stage, we used serial sections from the inferior temporal gyrus of a case with a neuropathological diagnosis of Alzheimer’s disease (Braak stage 5) and diffuse neocortical Lewy Body disease. The case had a PMI of 12 hours and was fixed for 72 hours. Immunostaining protocol steps were similar to those described for stage 1 up through the incubation of primary antibodies. Table 2 provides information on the primary antibodies. Notably, we included the MC1 antibody against conformationally altered tau proteins because it labels highly aggregated inclusions (Eckermann et al., 2007).

#### 2.3.2 Cycle of staining and stripping

After incubating the slides in a cocktail of the three primary antibodies, slides were washed thoroughly in PBST then incubated in 1:400 goat-anti-guinea pig IgG (H+L) HRP conjugated-secondary antibody (R-05076, Advansta) diluted in PBST for 60 minutes. Following additional PBST washes, the reaction was developed with TSA following the manufacturer’s instructions with a solution of 1:100 AF647 Tyramide (B40958, Thermo Fisher) and 1:100 of 100x H_2_O_2_ in 1x tris-buffered saline. The reaction was stopped with a PBST wash.

Following PBST washes and a transfer through 70% ethanol, the sections were treated with a solution of 0.8% Sudan Black-B in 70% ethanol for 35 minutes to block for lipofuscin. After 35 minutes, two 10-second washes in 70% ethanol was used to remove excess Sudan black-B. Sections were incubated in Sudan black-B between TSA development of NeuN and the fluorescent-conjugated secondary antibodies to be consistent with the subsequent test rounds that omitted NeuN incubation and development.

After rehydration in PBS, sections were incubated in a cocktail of 1:400 Goat anti-Rabbit IgG (H+L) (A-11008, Life Technologies) and 1:400 AF546 Goat anti-Mouse IgG (H+L) (A-11003, Life Technologies) in PBS for 90 minutes. Following washing in PBS, the slides were coverslipped with Prolong Glass Antifade Mountant with NucBlue (P36981, Thermo Fisher) then imaged at 10x magnification. After imaging, the slides were submerged in PBST overnight and agitated in order to remove the coverslip. Sections were washed in additional PBST to remove any excess mounting media.

BME stripping solution was prepared according to Gendusa et al. (2014). Under a fume hood, 20ml of 10% SDS was mixed with 12.5ml of 0.5M Tris-HCl (pH 6.8), 67.5ml of deionized (Millipore) water, and 0.8ml of BME. The solution was heated to 56^ω^C before use. Sections were incubated in the heated BME solution for 30 minutes. Sections were washed in four 15-minute dH_2_O immersions and then washed in TBST for five minutes, followed by immersion in 5% milk PBS solution for 30 minutes. One slide of each set was left in the milk solution overnight. All other slides were re-incubated with the original primary antibodies of each set (except NeuN) followed by the appropriate incubation in 1:400 Goat anti-Rabbit IgG (H+L) and 1:400 AF546 Goat anti-Mouse IgG (H+L) for 90 minutes, Sudan black treatment, mounting and imaging. The procedure was repeated up to 3 more times (Fig 1).

#### 2.3.3 Imaging acquisition

See the process described for Stage 1 (see 2.2.4)

## 3. Results

### 3.1 Stage 1: Evaluation of stripping techniques

#### 3.1.1 Phospho-tau antibody CP13

Only the heat-based and BME-based stripping methods were efficient in stripping this antibody, as noted by the lack of AF546 signal (Fig. 2A & B). However, tissue dissociated from the slide after the heat treatment. AF546 signal overlapping with AF647 signal on slides treated with glycine/SDS at low pH-based and commercial denaturing solution-based stripping techniques indicated a partial stripping of the antibodies (Figure 2C & D). In all stripping techniques, colocalization between AF488 and AF647 signals was nearly complete, indicating that antigenicity of the phospho-Ser202 Tau epitope detected by CP13 was preserved through the one round of stripping. Curiously, we noted higher AF488 than AF647 signal in the slides treated with heat-based and glycine/SDS at low pH-based methods, suggesting that these two strategies may expose additional epitopes for antibody reactivity. Table 3 summarizes these results

**Table 3:** Summary of outcomes of various stripping techniques from Stage 1.

| Strategy | Effective stripping |  |  | Antigen preservation |  |  | Tissue preservation |  |  |
| --- | --- | --- | --- | --- | --- | --- | --- | --- | --- |
|  | CP13 | NeuN | GFAP | CP13 | NeuN | GFAP | CP13 | NeuN | GFAP |
| Heat | + | + | + | + | + | + | - | - | - |
| Glycine/SDS at low pH | - | + | + | + | + | + | + | - | - |
| Denaturing solution | - | + | + | + | + | + | + | + | + |
| BME | + | + | + | + | + | + | + | + | + |

#### 3.1.2 Antibodies to label cell-types

All the stripping strategies were efficient in stripping antibodies, as shown by absence of AF546 signal (Fig. 3 and 4). Also, colocalization between AF488 and AF647 was nearly complete in all slides, indicating that the antigenicity of the epitopes for these antibodies was preserved after one round of stripping. In fact, higher AF488 than AF647 signal was noted in all strategies tested, suggesting that either the stripping reagents may expose additional epitopes for antibody reactivity, or, just reflect that most cameras are more sensitive to GFP than to Cy5. Once again, sections treated with heat-based method dissociated from slides. Table 3 depicts a summary of these results.

Overall, the BME stripping strategy was the only one meeting our criteria.

### 3.1 Stage 2 results

#### 3.1.1 Stripping efficacy

This was evaluated after each round by imaging the slide set aside as the stripping control (Fig. 5). These results were dichotomized as either ‘complete stripping’ or ‘incomplete stripping’, as quantitative evaluation of fluorescent signal can be imprecise. BME-based stripping achieved complete stripping of all antibodies tested, including MC1, through all rounds (Fig. 5).

**Figure 5:**
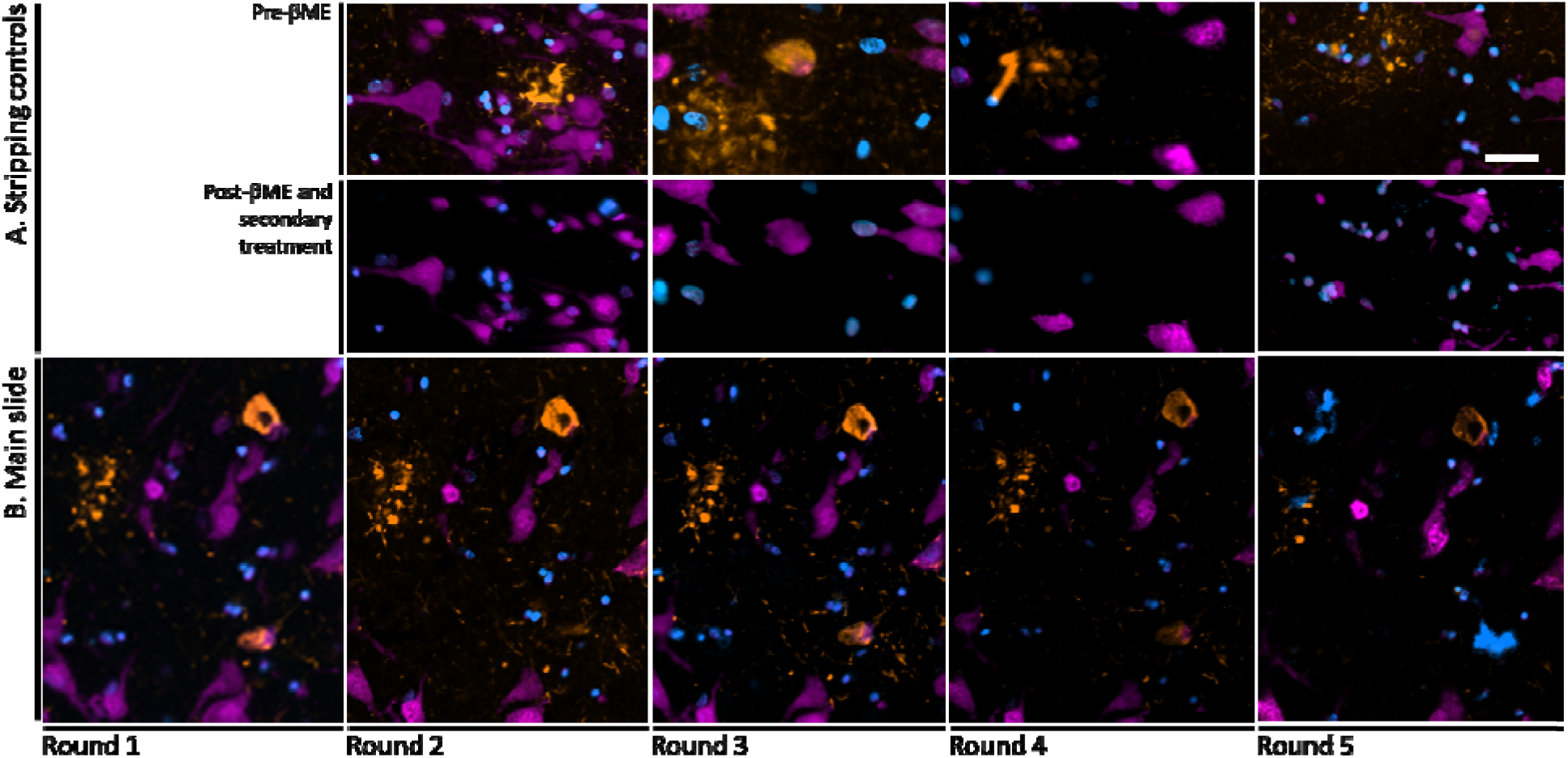
MC1 (orange) across five rounds of staining, stripping, and re-staining using βME-based stripping along with TSA-developed NeuN (pink) and Hoechst 33342 (blue). (A) Stripping controls which were stained in each round up until the final elution for that section where stripping was confirmed by confirming the lack of signal after development with AF546-conjugated secondary. (B) Signal from MC1 (orange) following staining, stripping, and re-staining across five rounds. NeuN (pink) was only developed in round 1 with AF647-TSA. Hoechst 33342 was stained in each round, as it is in the Prolong mounting media. Signal from MC1 appears across each of the rounds in the same locations. However a decrease in signal intensity is noted by Round 5.. Scale bar: 10 μm

#### 3.1.2 Tissue integrity

For all antibody sets tested, tissue sections had begun to dissociate from the glass by the fourth round. Thus, for the experimental conditions described here, four rounds of staining are viable, indicating that up to 11 antibodies could be probed in the same section of post-mortem human brain tissue.

#### 3.1.3 Evaluating of antigen preservation

Antigenicity was preserved throughout all the four rounds in which tissue remained intact, indicating that BME has little effect on altering epitope conformation, at least for the targets checked. Although we only used NeuN in round 1, the TSA-deposited AF647, marking NeuN, qualitatively maintained the original signal strength throughout all rounds (Fig. 5). Similarly, Hoechst 33342 signal was noted in each round after coverslipping using Prolong with NucBlue (Fig. 5).

#### 3.1.4 Composite image

To further validate this pipeline in practice, we applied this same protocol to a single section but with different antibodies (all listed in Table 2) in each round of staining, using TSA-deposited AF647 to mark NeuN in the first round and Hoechst33342 to co-register images (Fig. 6).

**Figure 6:**
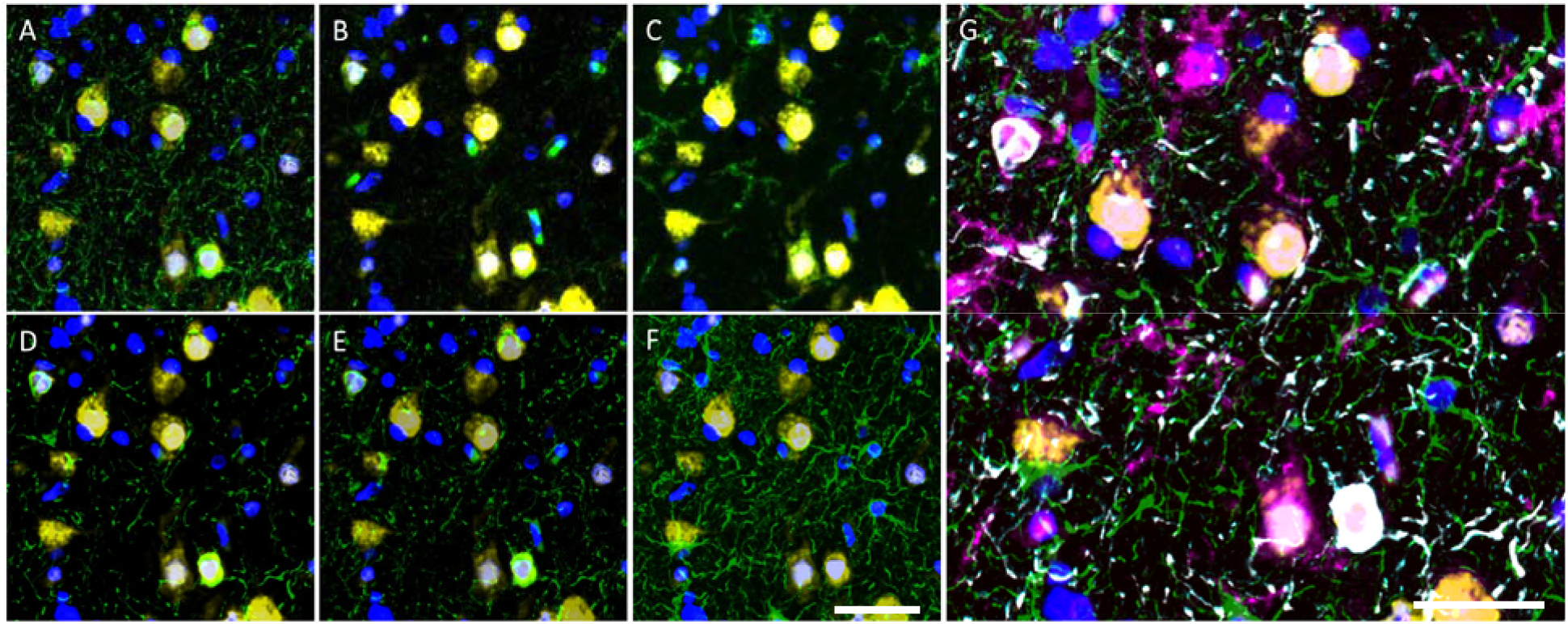
Multi-round immunofluorescence from three rounds of staining, separated each by a BME elution. Each round included an anti-Ms and anti-Rb primary antibody and a stripping control. Round 1 also included NeuN (yellow) developed with AF647. Each round was coverslipped with Hoechst33342 (blue). In panels A-G, NeuN is yellow and Hoechst33342 is blue. In green, panel A has CP13, B has MC1, C has TMEM119, D has T18, E has PHF-1, and F has GFAP. In panel G, blue is Hoechst33342, green is GFAP, pink is TMEM119, yellow is NeuN, red is MC1, gray is T18, cyan is PHF-1, and CP13 is orange. Because of the high overlap between tau inclusions, CP13, MC1, T18, and PHF-1 cannot be distinguished in the composite and appear as white.. Scale bar: 10 μm

## 4. Discussion

In this study aiming to establish a standardized protocol for multiplex immunofluorescence in postmortem human brain tissue for investigating neurodegenerative diseases with the resources of standard histopathology labs, we examined the efficacy of several previously reported techniques for antibody stripping. We focused on the antibody stripping step to simultaneously address limitations imposed by the restricted number of detection channels available in standard fluorescence microscopy and by possible cross-linking when using conspecific antibodies. From the seven main stripping strategies identified in the literature, we found that BME-induced stripping was the only one to enable complete stripping of all antibodies tested, including MC1 which binds to highly aggregated tau inclusions, while antigen and tissue integrity remained preserved across multiple rounds. In our hands, microwave heating of samples in citrate buffer for antibody stripping, the recommended strategy for stripping antibodies in TSA-based multiplex immunostaining protocols from ThermoFisher Scientific (Company publication #MAN0015834) and PerkinElmer (“Opal Multiplex IHC Assay Development Guide”), caused early tissue dissociation from the slide.

In Stage 2, we tested tissue and antigen integrity across five rounds of BME-induced stripping in a variety of antibodies. MC1 was of particular interest, as it targets a conformation-specific epitope of tau known to occur in neurofibrillary tangles. The potential for the stripping technique to alter the target confirmation was of concern, yet the signal from MC1 remained robust throughout each of the five rounds in Stage 2 (Fig. 5). Additionally, we show that the use of Hoechst 33342 in each round of staining or TSA reporting of a neuronal target in the first round of staining are good options as landmarks for co-registration of images in different rounds is feasible with BME.

This pipeline becomes a valuable framework for use in standard labs lacking sophisticated equipment, particularly when incorporating open-source or proprietary imaging analysis tools, (Alegro et al., 2017; Camp et al., 2002; Marrero et al., 2016; McCabe et al., 2005). Although our protocol accommodates up to 11 markers, this number could be possibly increased in labs with access to fluorescent microscopes equipped with more than four channels.

Although we implemented rather than created a new approach for multiplex staining, our protocol was optimized for the use with human postmortem brain tissue, which is more prone to dissociation, and antibodies targeting neurodegenerative inclusions, which requires a more potent stripping method. Thus, we are better suited to evaluate different stripping techniques that were previously suggested to work universally, applying techniques used in both immunohistochemistry and western blotting. While only testing a limited number of antibodies, we provide an easy strategy to evaluate if a stripping strategy is effective for a given antibody and at the same time test if the stripping technique affects antigen integrity for designing and validating multiplex immunostaining protocols. By utilizing TSA in stage 1, our reporter (signal from AF546) is extremely sensitive to residual antibodies left after stripping steps that might otherwise be missed if using traditional immunofluorescence (Chao et al., 1996; Wang et al., 1999).

In conclusion, we presented a multiplex immunostaining protocol that can be easily implemented in standard labs and is appropriate for use in suboptimal tissue and can incorporate a large array of antibodies. We detailed the protocols and the proposed quality control steps to validate antibody stripping. With proper validation, optimization, and implementation of controls, use of multiplex immunostaining protocols is realistic in standard histopathology labs and will likely be key for elucidating the molecular fundamentals of neurodegenerative diseases.

## 5. Acknowledgments

The authors thank the donors to the Neurodegenerative Disease Brain Bank of the University of California, San Francisco Memory and Aging Center and their families for their support of this work. The authors also thank Celica Cosme, Rana Eser, and Mackenzie Hepker for their technical assistance and laboratory support.

